# *Pseudomonas aeruginosa*-infected Myeloid-derived suppressor cells (MDSC) down-regulate lymphocyte activity and improves mice survival, following *in vivo* lung transfer

**DOI:** 10.1101/2024.04.24.590871

**Authors:** Maëlys Born-Bony, Clémentine Cornu, Bérengère Villeret, Romé Voulhoux, Jean-Michel Sallenave

**Affiliations:** Laboratoire d’Excellence Inflamex, Institut National de la Santé et de la Recherche Médicale, Physiopathologie et Épidémiologie des Maladies Respiratoires, Université Paris-Cité, Paris, France.; Laboratoire de Chimie Bactérienne LCB-UMR7283, CNRS, Aix Marseille Université, IMM, Marseille, France.

## Abstract

*Pseudomonas aeruginosa* (*P.a*.) is a pathogenic opportunistic bacterium, classified as a priority by the WHO for the research of new treatments. As this bacterium is harmful trough the inflammation and tissue damage it causes, we investigated the role of Myeloid Derived Suppressor Cells (MDSC) in *P.a.* infections and their potential as a therapeutic target. We found that upon *P.a.* exposure, MDSC activity is increased and gain contact-independent properties. Interestingly, this activation is dependent on *P.a.* mobility but not its flagellin nor TLR5-MyD88 pathway. We also show that MDSC adoptive transfer increases mice survival in *P.a.* acute lung infection both in therapeutic and prophylactic set ups. Finally, using *an in vitro* scratch assay model, we suggest that MDSC acts directly on lung epithelium to stimulate its repair. Together, we highlight a potential beneficial role of MDSC in *P.a.* infection response. We believe that the unique properties of MDSC make them attractive potential new therapeutic tools for patients with acute or chronic inflammatory diseases, where inflammation has to be kept in check.

## INTRODUCTION

*Pseudomonas aeruginosa* (*P.a*.) is an opportunistic bacterium causing airway infections particularly in cystic fibrosis (CF) and intensive care units (ICU) ventilated patients. Infection by this pathogen leads to airway inflammation, severe lung tissue damage and can result in respiratory insufficiency and death in up to 40% of cases^1–3^. *P.a*. shows high antibiotics resistance and is classified by the World Health Organization (WHO) as one of the priority pathogens for the research of new treatments^4^.

Our group has previously shown that IL-6-elafin (the latter being an anti-inflammatory/regulatory molecule)^5,6^ genetically modified macrophages adopted a regulatory profile and improved mice survival when used as prophylaxis in a *P.a*. infection model^7^. These results suggested that diminishing inflammation in *P.a*. infection is key to reduce lung damage and resolve the infection. Other studies also demonstrated that adoptive transfer of immune cells such as macrophages is beneficial for *P.a*. elimination from the lung, resolution of inflammation and overall outcome^8,9^.

Important regulatory cells of the immune system are myeloid derived suppressor cells (MDSC). MDSC constitute an immature heterogeneous cell population with strong immunosuppressive effect on T cell responses, accompanied with Treg and regulatory macrophages induction^10,11^. They are constituted of two major sub-populations: polymorphonuclear (PMN) and monocytic (M)–MDSC. These cells are absent at steady state but their proportion increases dramatically in pathological condition such as cancer, or in some infectious diseases^12^. Importantly, depending on the pathogen involved, MDSC presence can either be detrimental or beneficial. For example, MDSC have been described to be responsible of excessive immune suppression in *Mycobacterium tuberculosis* infection^13^. By contrast, MDSC have been associated with reparative functions and an ameliorated prognosis in *Klebsellia pneumoniae* or *Candida albicans* infections.^14–18^.

Regarding *P.a*., Rieber et al. showed that MDSC proportion in total PBMC is increased in CF patients infected with *P.a* and that the presence and proportion of these cells correlate positively with ameliorated lung function^19^. In mice, the same group showed recruitment of MDSC after *P.a* infection, but their role and mode of action remained unspecified^20^.

Here, we evaluated how MDSC function is modulated after *P.a in vitro* infection and whether these cells could be used in adoptive transfer in a therapeutic setting. We first describe a new model of *in vitro* (ER-Hoxb8) differentiated MDSC, and show that these MDSC are functionally activated and gain contact-independent regulatory properties upon *in vitro P.*a infection (T cells proliferation inhibition and epithelial repair induction). Crucially, we also demonstrate that these cells adoptively transferred in a therapeutic setting improved lung inflammation in *P.a* infected mice.

## RESULTS

### MDSC can be differentiated *in vitro* from fresh mice bone marrow or ER-Hoxb8 cells

Although MDSC cannot be isolated from healthy mice, it is possible to induce their differentiation from bone marrow in a media containing IL-6 and GM-CSF^21^. In addition to these primary cells, we also used a strategy of differentiation from immortalized ER-Hoxb8 cells, which are derived from C57Bl/6 mice bone marrow progenitors transduced by an estrogen receptor – Hoxb8 fused construction^22^.

Using the above-mentioned IL-6 and GM-CSF differentiation protocol (Fig. 1A), we induced BM- and ER-Hoxb8-derived MDSC and showed that they expressed classical markers of MDSC : CD11b+ CD11c-Ly6C+, Ly6G+ for PMN-MDSC and Ly6G-for M-MDSC^11^ (Fig. 1B and C). These two populations were also distinguishable when cells were imaged after cytospin (Fig. 1D). Although BM-MDSC and ER-Hoxb8 expressed similar markers with similar sub-population repartition, ER-Hoxb8 MDSC expansion was greater than the one achieved for BM-MDSC, revealing an important benefit of using this model (Fig. 1E).

**Figure 1.**
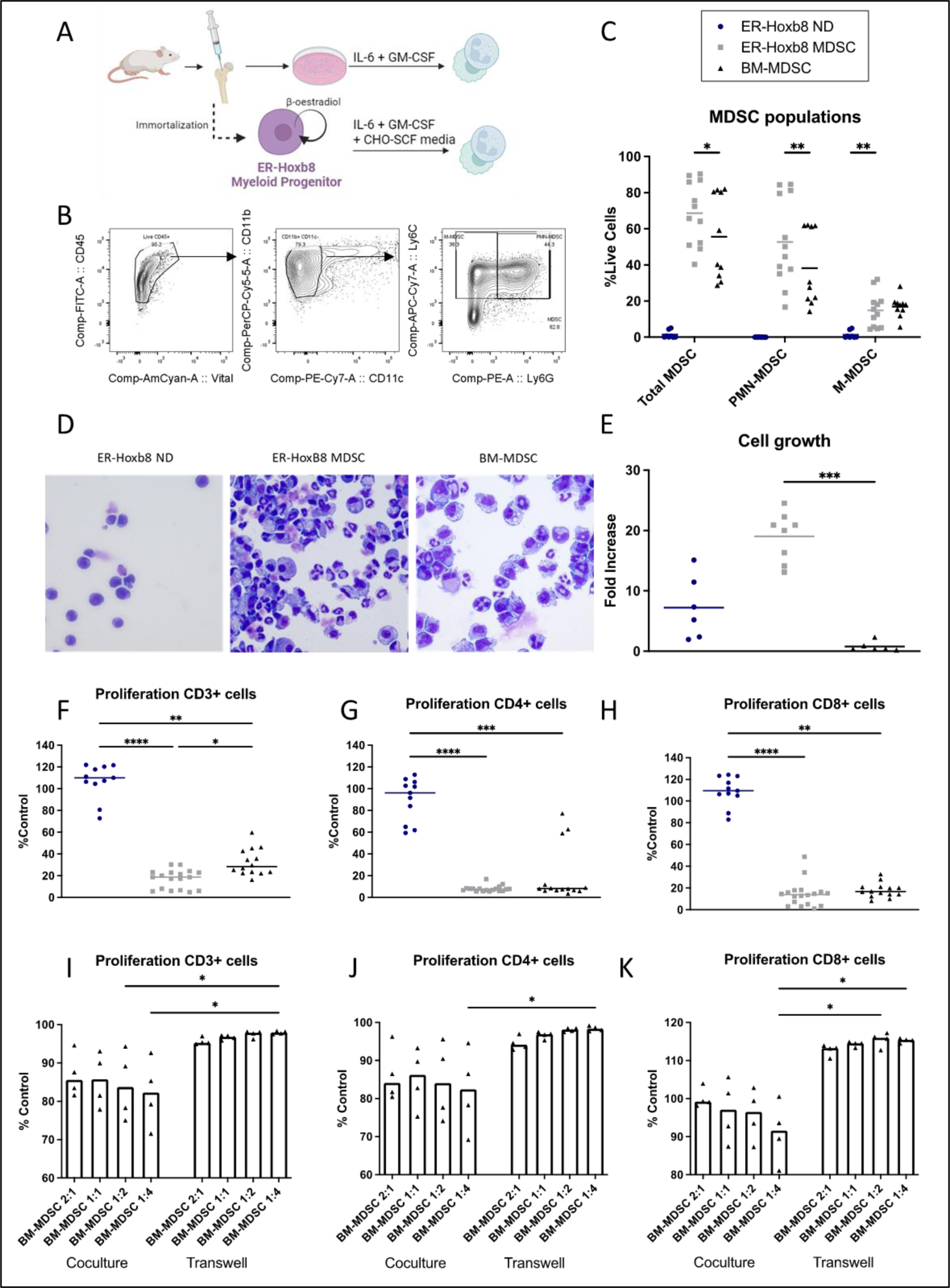
Cells differentiated from bone marrow and ER-Hoxb8 are fully functional MDSCs and require cell contact for their T cell-inhibitory activity. MDSCs were differentiated from C57Bl6J mice bone marrow cells (BM-MDSC) in presence of IL-6 and GM-CSF or from ER-Hoxb8 cell line (ER-Hoxb8 MDSC) in presence of IL-6 and GM-CSF and CHO-SCF media for 5 days (A). Cells were characterized by flow cytometry (B-C) and cytospin (D). Flow cytometry gating strategy (B) is shown on BM-MDSC. Cell growth of BM- and ER-Hoxb8-MDSC was calculated as number of cells after differentiation / number of cells seeded at D0 (E). Inhibitory capacity of BM- and ER-Hoxb8-MDSC on T cells was assessed in a co-culture proliferation assay (1:1 ratio). T cells were stained with Cell Trace Violet (CTV) and stimulated for proliferation with CD3/CD28 Dynabeads for 72h. CTV dilution was measured by flow cytometry; results are expressed as %Control (%proliferation sample/% proliferation of T cell only) (F-H). Inhibitory capacity of BM-MDSC was also evaluated in a direct co-culture or Co-Star Transwell plate in the same condition as before (I-K). Data show median values (n = 5 independent experiments, 1 to 5 replicate per experiment). Statistical significance: Kruskal-Wallis, Dunn’s multiple comparison, ∗p < 0.05; ∗∗p < 0.01; ∗∗∗p < 0.001; ∗∗∗∗p < 0.0001.

Since marker expression is not sufficient for MDSC identification, we next evaluated MDSC function in a T cell proliferation assay. MDSC were cocultured with T cells at 1:1 ratio and the latter were stimulated for proliferation with CD3/CD28 Dynabeads in a media containing IL-2. T cells cultured with BM or ER-Hoxb8 MDSC, but not with non-differentiated ER-Hoxb8 (ER-Hoxb8 ND), showed inhibited proliferation only when cells were in direct contact (Fig. 1F-H), but not when they were separated in a Transwell cell culture system (Fig. 1I-K). Overall, this demonstrated that fully functional MDSC can be produced from either bone marrow cells or from the ER-Hoxb8 cell line and that MDSC have a contact dependent inhibition of proliferation.

### PAO1 infection increases MDSC inhibitory activity

To assess cytokine production, BM-MDSC were infected with the *P.a* PAO1 laboratory strain for 6 hours, and gene expression and supernatant cytokine output was then evaluated by qPCR and ELISA. (Fig. 2A). Even though the protein (and genes coding) for S100A8/9, involved in MDSC proliferation and accumulation, was not impacted by infection (Fig. 2B, F, J), infected MDSC showed increased expression of genes associated with MDSC functionality such as iNos, Nox1 (Fig. 2C, E). IL-6, IL-10. LCN2 levels were also increased at the protein level as measured by ELISA (Fig. K-N). PD-L1 expression was increased at the RNA levels (Fig. 2O), as were the soluble and membrane forms of the protein (Fig. 2P-R).

**Figure 2.**
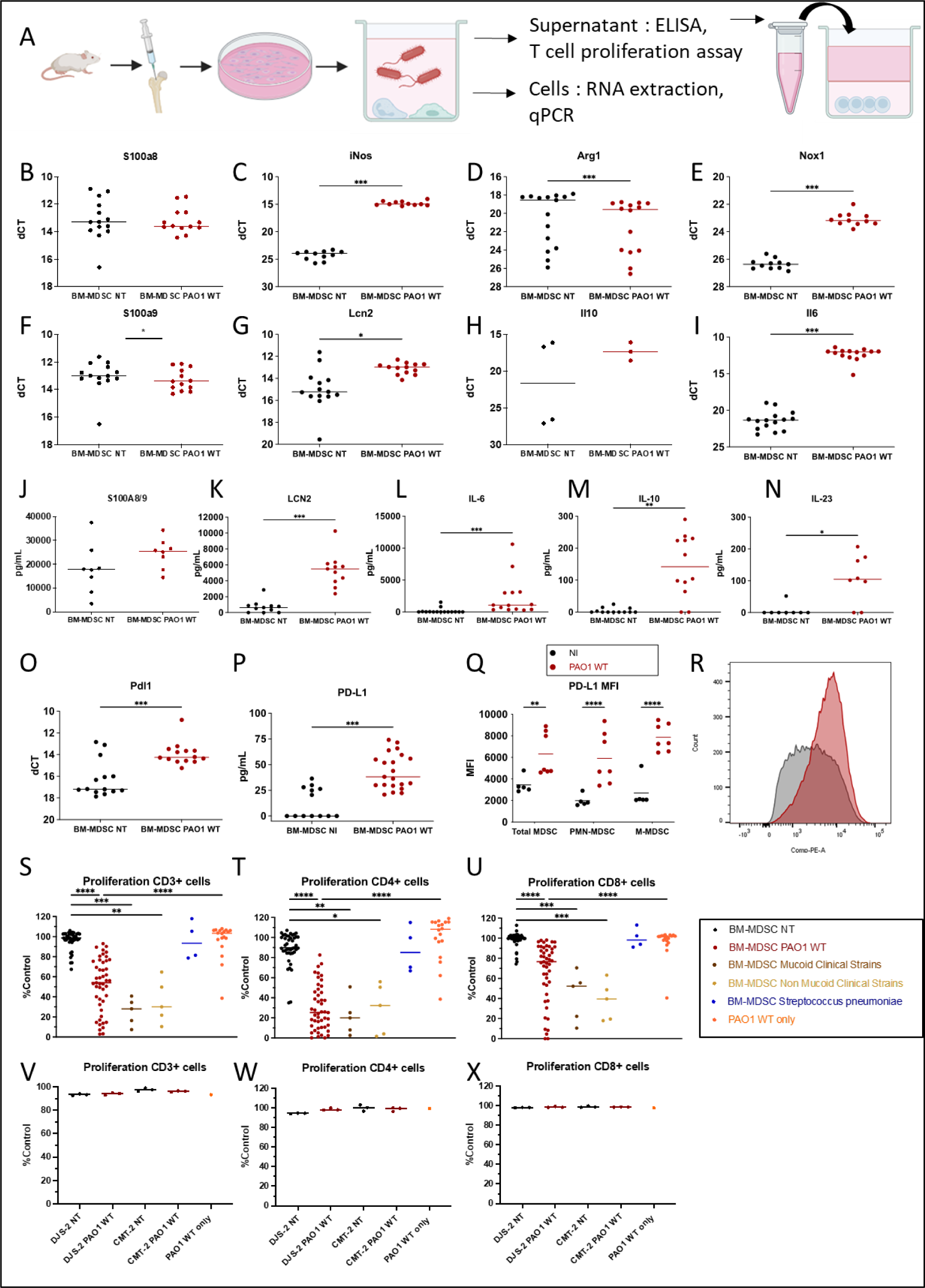
Infection of BM-MDSC with *Pseudomonas aeruginosa* up-regulates inflammatory markers and unleashes their T-cell inhibitory activity. *In vitro* differentiated BM-MDSCs were mock-(NT = non-treated) or infected with PAO1 WT *P.a* strain, *P.a* clinical strains or *Streptococcus pneumoniae (S.p)* for 6 hours at MOI 1. MDSCs were retrieved and lysed for RNA extraction and qPCR analysis with the dCt method (B-I, O) was performed, with dCT = CT gene of interest - CT housekeeping gene (18S). In some experiments, cells were harvested and stained for PD-L1 and MDSC subset markers by flow cytometry (Q, R). Data show median values (n = 3 independent experiments, 1 to 5 replicate per experiment). Statistical significance: Wilcoxon, ∗p < 0.05; ∗∗p < 0.01; ∗∗∗p < 0.001; ∗∗∗∗p < 0.0001. Cell supernatants were also retrieved, and cytokine protein concentration in measured by ELISA (J-N, P). For some experiments (panels S-U), supernatant of PAO1 WT alone (PAO1 only) was also retrieved. T cells were exposed to filtered supernatants and their proliferation was evaluated in a 72 hours T cell proliferation assay, using T cell Cell Trace Violet dilution as a proliferative index (flow cytometry, panels S-U)). For the “BM-MDSC NT” and “BM-MDSC PAO1 WT” groups, data from all the experiments realized in this study (Fig. 2 and 5) are pooled (n>10). For the other groups, data come from n=3 independent experiments, 1 to 2 replicate per experiment, which are only presented in this figure. Statistical significance: Kruskal-Wallis, Dunn’s multiple comparison, ∗p < 0.05; ∗∗p < 0.01; ∗∗∗p < 0.001; ∗∗∗∗p < 0.0001. Finally, epithelial cells lines CMT-2 (alveolar) and DJS-2 (Club cells) were infected with PAO1 WT as for MDSCs (V-X). Supernatant was retrieved and filtered before being used in a T cells proliferation assay. Data show median values (n = 1 independent experiments, 3 replicates per experiment).

Filtered supernatants from BM-MDSC-infected bacteria were then added to T cells cultures and T cell proliferation was then measured. We found that CD3+, CD4+ and CD8+ T cell proliferation was inhibited by exposure to infected MDSC supernatant but not by non-treated (NT) MDSC (Fig. 2S-U). Exposition to PAO1 only (no MDSC) supernatant was used as control and showed no inhibition of T cell proliferation. Importantly, this T cell proliferation inhibitory activity was not restricted to MDSC infected with the PAO1 strain but was also evident in cells infected with clinical isolates from patients with CF, irrespective of their mucoid status (Fig. 2S-U). Equally noteworthy was the demonstration that this activity was not present in MDSC infected with the Gram+ *Streptococcus pneumoniae*, another clinically important lung pathogen (Fig. 2S-U).

To assess whether the observed effect was specific to MDSC, we reproduced the infection experiment this time on epithelial cells. CMT-2 (alveolar) or DJS-2 (Club) cell line were infected with PAO1 for 6 hours at MOI 1 and their supernatant retrieved for T cell proliferation assay (Fig. 2V-X). Supernatant of epithelial cells did not induce T cell proliferation inhibition indicating some specificity of the previously observed effect.

Infection experiments were reproduced in the ER-Hoxb8 MDSC model (Fig. 3). ER-Hoxb8 displayed similar properties as BM-MDSC at steady sate and after infection. Gene expression evaluated by qPCR showed comparable expression of *iNos*, *Nox1*, *S100A8*, *S100A9*. Expression and secretion were also similar in both models for Lcn2, IL-10 and PD-L1. The only differences observed between the two models were higher Arg1 expression (Fig. 3B) in ER-Hoxb8 MDSC when compared to BM-MDSC both at steady state (called mock- or NT = not treated in the figures) and after infection, and reduced IL-6 secretion in ER-Hoxb8 MDSC after infection (Fig. 3H). Importantly, these slight differences did not affect MDSC inhibitory activity post infection, as it was similar in both models (Fig. 3L-N, compared to Fig 2S-U), demonstrating the use and validity of both set-ups for our study.

**Figure 3.**
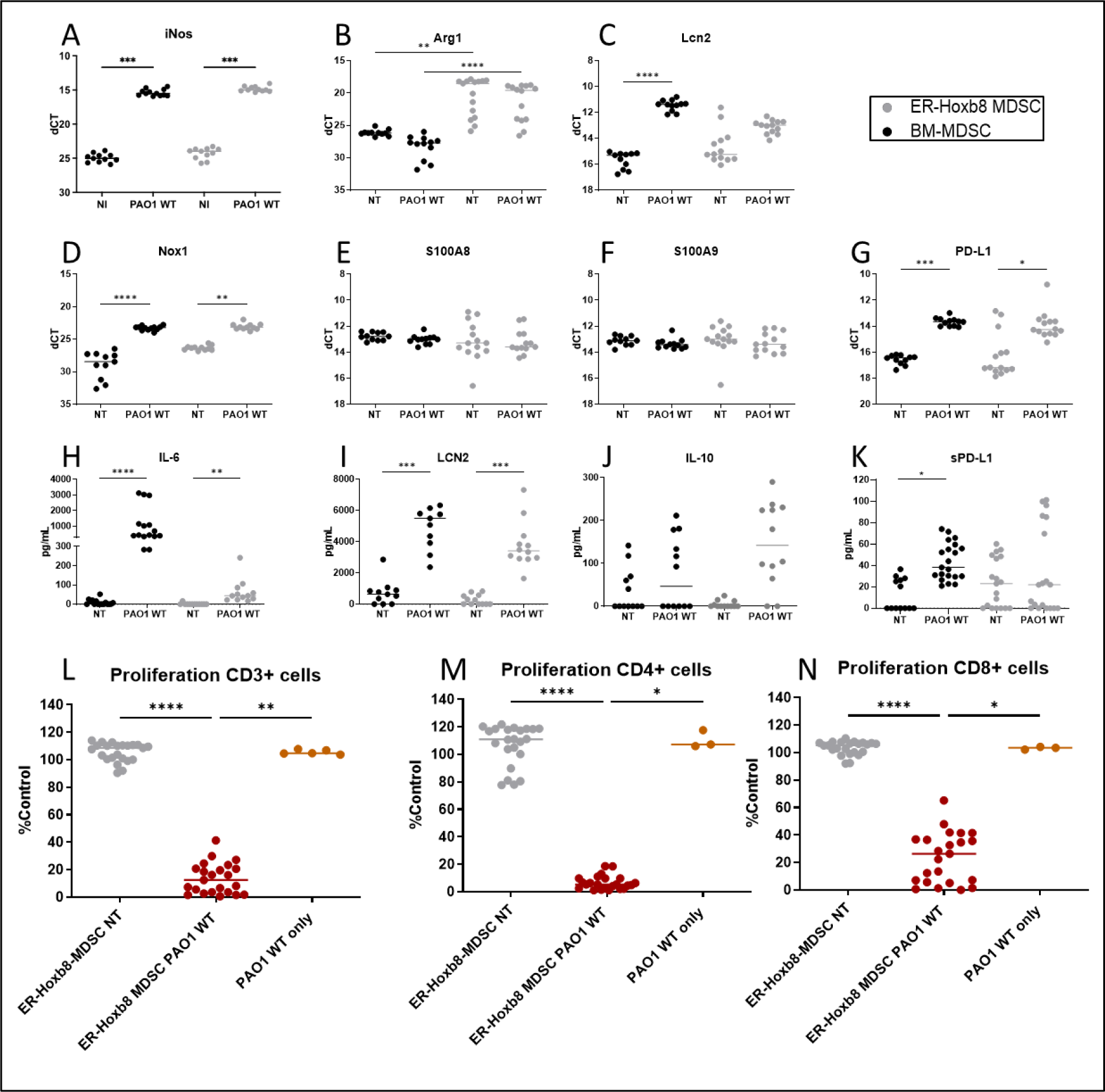
Infection of ER-Hoxb8-MDSCs with *P.a* up-regulates inflammatory markers and uncovers their T-cell inhibitory activity. *In vitro* differentiated ER-Hoxb8**-**MDSCs were mock-(NT) or infected with PAO1 WT for 6 hours at MOI 1. Cells were retrieved and lysed for RNA extraction and qPCR analysis with the dCt method (B-G). Supernatants were collected and filtered on 0.2µm filters. Protein concentration in supernatant was measured by ELISA (H-K). Data show median values (n = 3 independent experiments, 1 to 5 replicate per experiment). BM-MDSC data points were pooled from all experiments to enable comparison with ER-Hoxb8 MDSCs and thus are also represented in figure 2. Statistical significance: Wilcoxon, ∗p < 0.05; ∗∗p < 0.01; ∗∗∗p < 0.001; ∗∗∗∗p < 0.0001. Inhibitory capacity of MDSC supernatant was evaluated in a 72 hours T cell proliferation assay. T cells proliferation was assessed through Cell Trace Violet dilution measurement via flow cytometry (L-N). For some experiments (panels L-N), T cells were also treated with supernatants of PAO1 (PAO1 WT only). Data show median (n = 5 independent experiments, 1 to 5 replicate per experiment). Statistical significance: Kruskal-Wallis, Dunn’s multiple comparison, ∗p < 0.05; ∗∗p < 0.01; ∗∗∗p < 0.001; ∗∗∗∗p < 0.0001.

### Both PMN-MDSC and M-MDSC show increased inhibitory activity after PAO1 infection

Since MDSC constitute a heterogeneous population made of neutrophils (PMN) and monocytes (M), (Fig. 4B), we separated BM-MDSC by immunomagnetic techniques (see strategy in Fig. 4A) and analyzed them by FACS (Fig. 4B-E) and cytospin (Fig. 4F-I). After differentiation (Fig. 4B), they were magnetically separated in three groups: a) Ly6G high PMN-MDSC (Fig. 4C, G, composed of Ly6G+ Ly6C+ cells), b) positively selected GR-1+ cells, composed of a mix of LyC high Ly6G mid (Fig. 4D, H), and c) negatively selected GR-1-cells, composed of mostly Ly6C high Ly6G neg M-MDSC (Fig. 4E, I). Because the yield was too low in group b), only groups a) and c) were separately infected with PAO1 as per previous experiments. Although inhibition of CD3+ T cell proliferation was only statistically significant for the supernatant from group C cells (Fig. 4J), a trend was also present for infected Ly6G high PMN-MDSC (group A cells), suggesting that both myeloid cell types (neutrophils and monocytes) harbored this T cell inhibitory activity (Fig. 4J-L).

**Figure 4.**
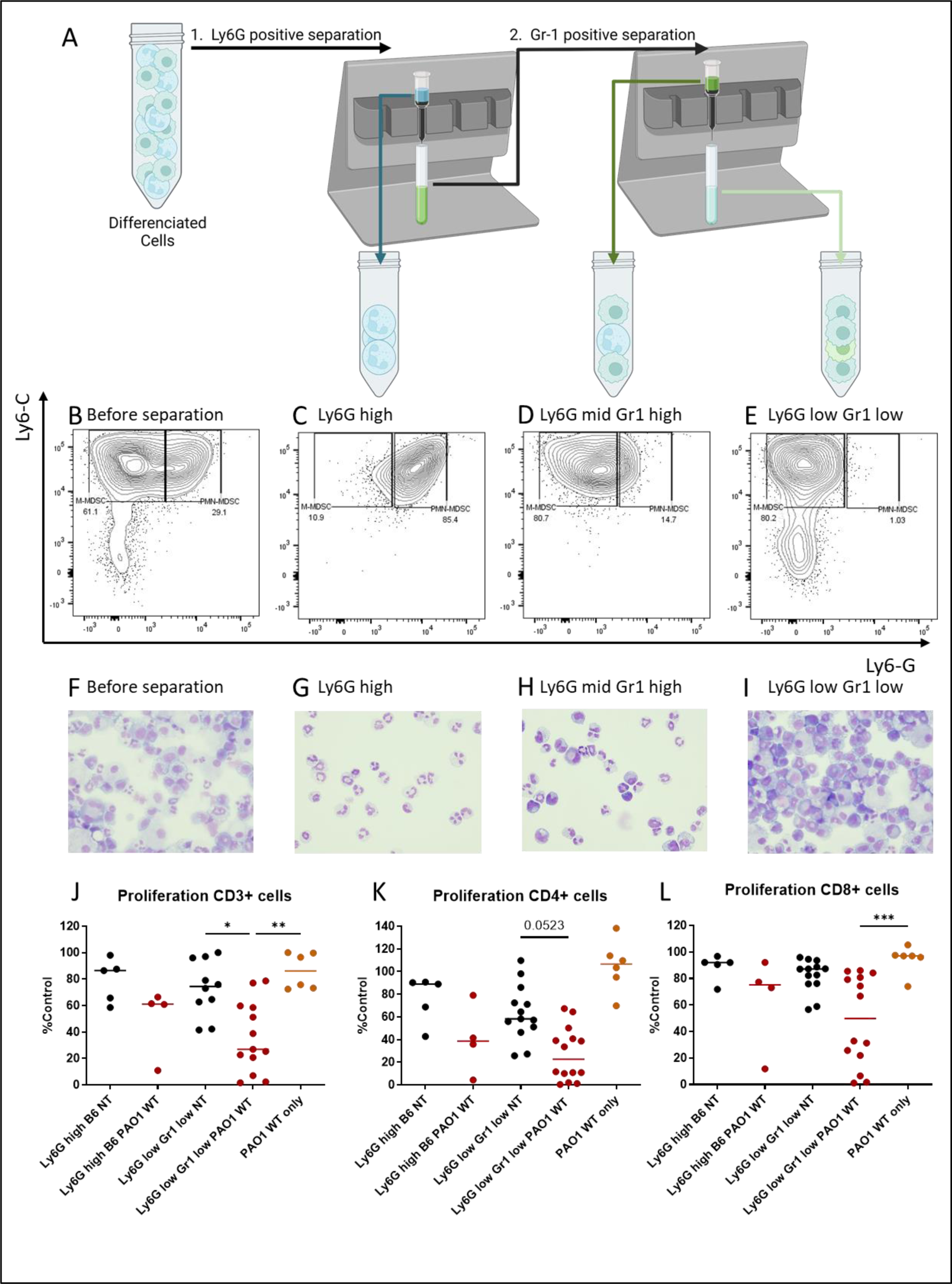
Immunomagnetic separations of BM-MDSCs fractions show that both PBMCs Ly6G high PBMCs and monocytes Ly6C high (Ly6G low Gr1 low) inhibit lymphocyte proliferation after PAO1 infection. *In vitro* differentiated MDSCs were magnetically separated in subpopulations according to their Ly6C/Ly6G expression (A) and were either mock-(NT) or infected with PAO1 WT for 6 hours at MOI 1. MDSC markers expression and morphology was assessed in different fractions by flow cytometry (B-E) and cytospin (F-I). Supernatants were collected and filtered on 0.2µm filter. Inhibitory capacity of MDSC supernatant was evaluated in a 72-hour T cell proliferation assay. Independently, T cells were also treated with supernatants of PAO1 (PAO1 WT only). T cells proliferation was assessed through Cell Trace Violet dilution measurement via flow cytometry (J-L). Data show median values (n = 4 independent experiments, 1 to 4 replicate per experiment). Statistical significance: Kruskal-Wallis, Dunn’s multiple comparison, ∗p < 0.05; ∗∗p < 0.01; ∗∗∗p < 0.001; ∗∗∗∗p < 0.0001.

### MDSC inhibitory activity requires live motile bacteria and is TLR independent

To further dissect the mechanisms involved in *P.*a-mediated MDSC modulation, we demonstrated that the latter was dependent on the presence of live bacteria since different doses of heat inactivated PAO1 were unable to confer MDSC with this inhibitory activity (Fig. 5A-C).

**Figure 5.**
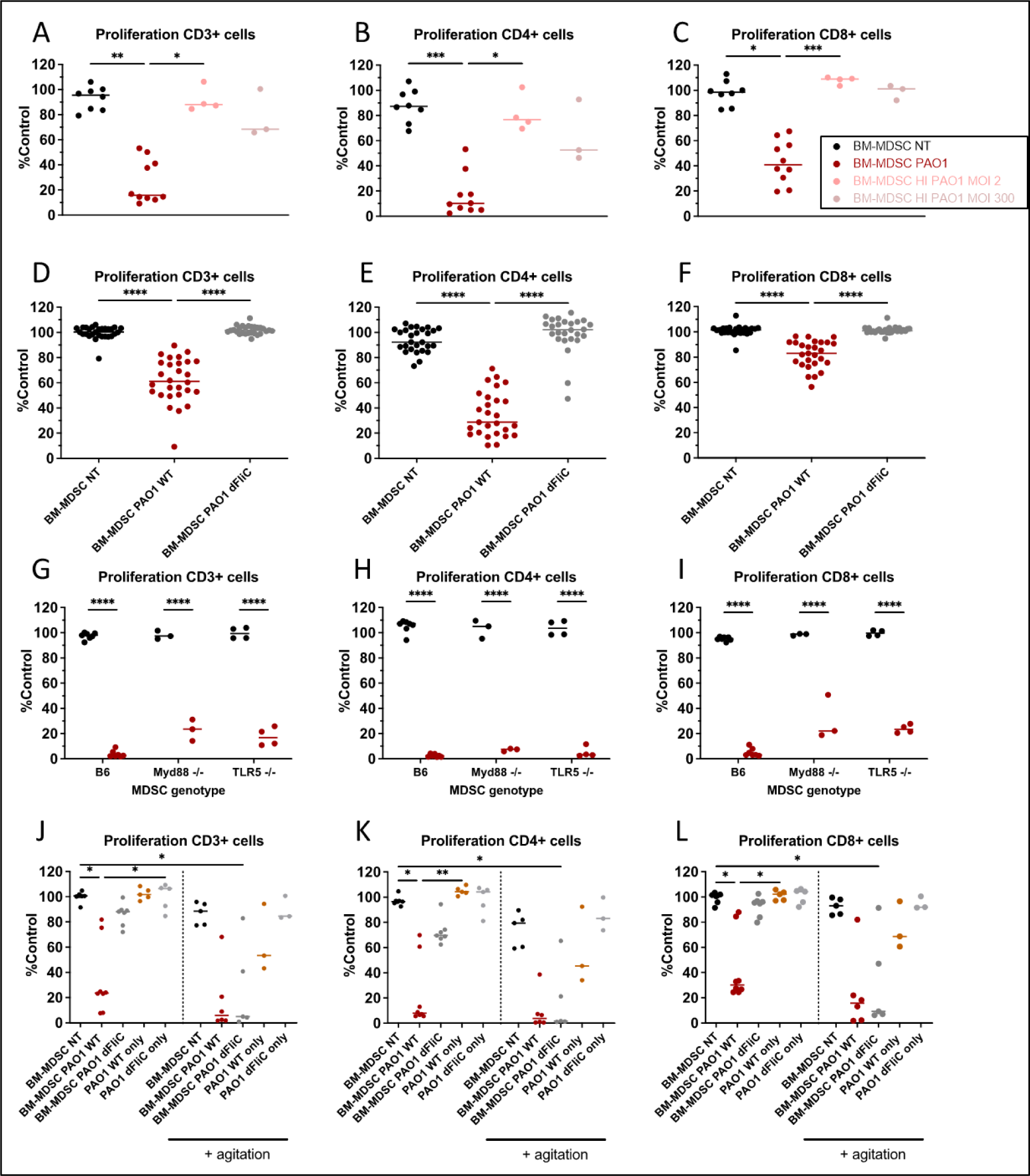
The induction of inhibition in MDSC supernatants requires live and mobile bacteria and is independent of the TLR/Myd88 pathway. *In vitro* differentiated C57/Bl6 WT (A-F), TLR5 or Myd88 knock-out (G-I) BM-MDSCs were mock-(NT) or infected with live or heat inactivated (HI) PAO1 (A-C) or dFliC PAO1 (D-F) for 6 hours at MOI 1. PAO1 dFliC infection was independently repeated on a plate with agitation (J-L). Supernatants were collected and filtered on 0.2µm filter. Inhibitory capacity of MDSC supernatant was evaluated in a 72-hour T cell proliferation assay. In subset experiments (panels J-L) T cells were also treated with supernatants of either PAO1 WT (PAO1 WT only) or PAO1dFliC (PAO1 dFLiC only) bacteria. T cells proliferation was assessed through Cell Trace Violet dilution measurement via flow cytometry. Data show median values (n = 3 independent experiments for A-C, G-L; n > 10 for D-F, with 1 to 5 replicate per experiment). Statistical significance: Kruskal-Wallis, Dunn’s multiple comparison, ∗p < 0.05; ∗∗p < 0.01; ∗∗∗p < 0.001; ∗∗∗∗p < 0.0001.

Furthermore, we tested the activity of important *P.a* virulence factors known to modulate myeloid activity^23–25^. Whereas d*lasA*, d*lasAlasB* (type 2 secretion mutants) and d*pscN* (a type 3 secretion mutant) were equally able to increase MDSC-mediated T cell inhibitory activity (not shown), the d*fliC* mutant (lacking the flagella) did not confer the cells with such activity (Fig. 5D-F). This suggested that flagellin (the protein monomer constituting the flagella) might be an important ligand for MDSC, but experiments adding purified flagellin to d*fliC* PAO1 did not restore the MDSC modulation (not shown). The fact that flagelin was likely not responsible for the described phenotype was further exemplified by experiments where BM-MDSC were generated from MyD88 -/- and TLR5 (the flagellin receptor) -/- mice and infected with PAO1. Indeed, these supernatants were still inhibitory (albeit slightly less, (Fig. 5G-I)), demonstrating that the PAO1-mediated MDSC modulation is mostly independent of the TLR-Myd88 pathway.

We then tested whether it was the reduced mobility of the d*fliC* mutant^26^ which hampered MDSC activity. To mimic bacteria mobility, MDSC were infected in a plate with agitation (Fig. 5J-L). In these conditions, MDSC infected with PAO1 d*fliC* tended to show the same inhibitory gain as PAO1 WT infected MDSC. These results indicate that neither flagellin nor bacterial mobility are necessary to induce MDSC contact-independent inhibitory activity.

### MDSC supernatant-T cell interaction

1) Heat- and Proteinase K-treatment do not abolish MDSC supernatants inhibitory activity

Investigating further ‘downstream’ mechanisms underlying the inhibitory activity of *P.a*-infected MDSC supernatants, we showed that the inhibitory activity was heat- and proteinase K-resistant (Fig. 6), suggesting that the inhibitory moiety(ies) might not be proteins.

**Figure 6.**
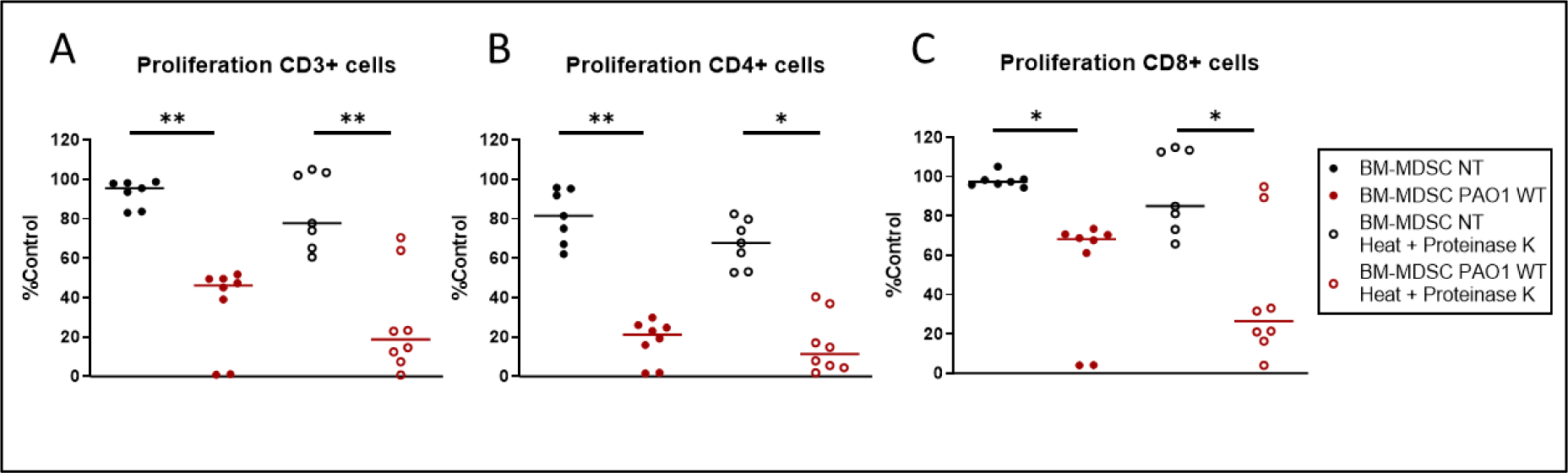
Inhibitory effectors in MDSC supernatant are resistant to heat and Proteinase K treatment. In vitro differentiated BM-MDSCs were mock- or infected with WT PAO1 for 6 hours at MOI 1 (A-C). Supernatants were collected, filtered on 0.2µm filters and treated for 1h at 56°C with 200ug/mL Proteinase K (PK). Heat treatment of supernatants for 30min at 95°C was performed in order to, in a single experiment, inactivate PK (to avoid spill over protease activity on T cells (see below)) and to test the heat resistance/sensitivity of the MDSCs-derived inhibitory factors. The inhibitory capacity of MDSC supernatants was evaluated in a 72-hour T cell proliferation assay. T cells proliferation was assessed through Cell Trace Violet dilution measurement via flow cytometry. Data show median value (n = 3 independent experiments, 2 to 3 replicate per experiment). Statistical significance: Kruskal-Wallis, Dunn’s multiple comparison, ∗p < 0.05; ∗∗p < 0.01; ∗∗∗p < 0.001; ∗∗∗∗p < 0.0001.

2) MDSC supernatants do not act solely at the level of the T cell synapse.

a) *CD3/CD28 stimulation*

To assess whether MDSC supernatant were acting at the synaptic level, or downstream of the T cells synapse, we used a different set up where activating beads were removed after 24 hours and proliferation let to proceed for a further 72 hours. MDSC supernatant were either incubated with the beads during the 24 hours, or, after the removal of the beads, during the 72 hours of T cells proliferation (Fig. 7A-C).

**Figure 7.**
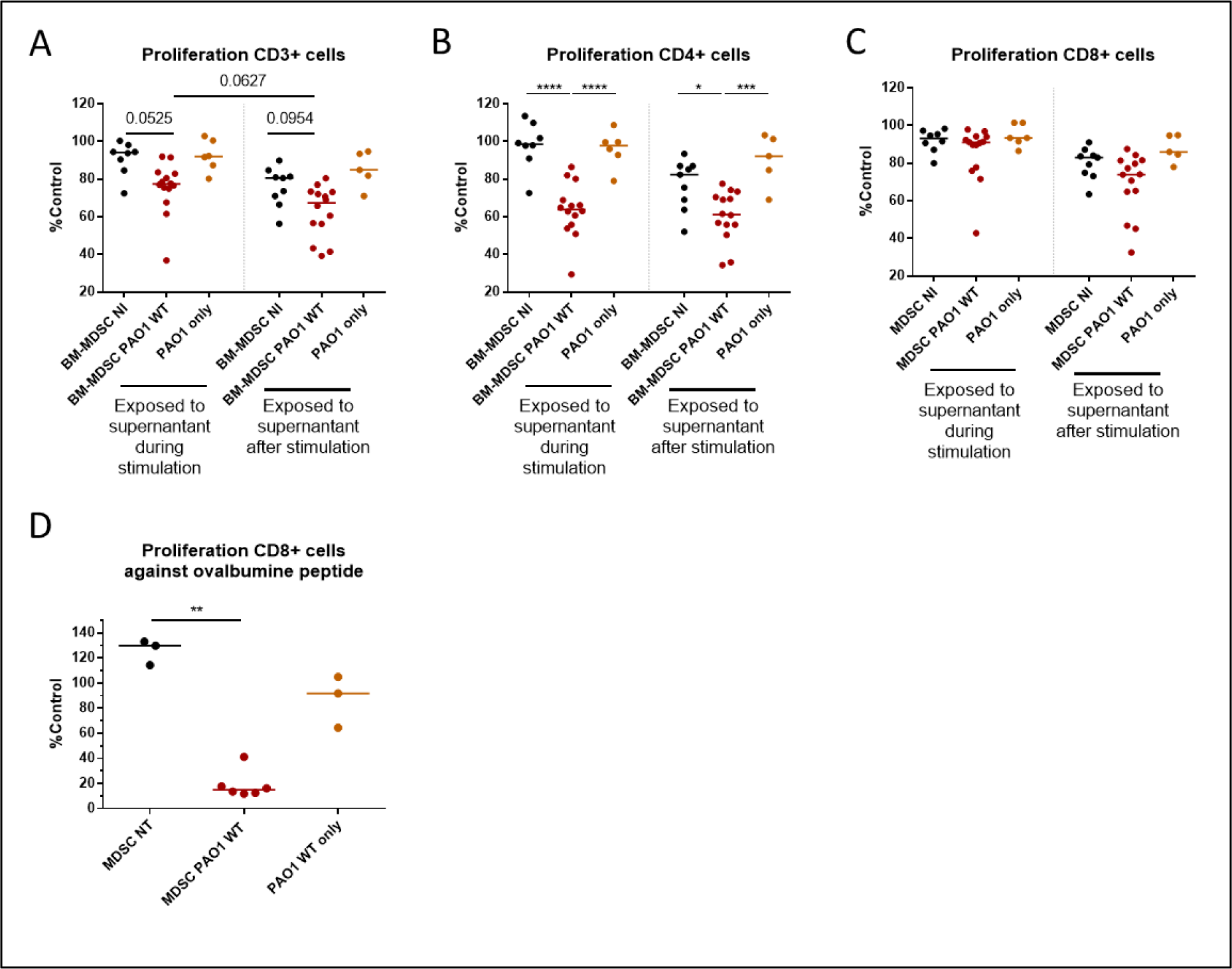
Study of MDSCs-T cells interactions with different stimulatory kinetics and with a specific agonist stimulus. Kinetics of T cells proliferation: T cells were exposed to activating beads for 24hours. Beads were the removed and T cells proliferation let to process for another 72 hours. T cells were either exposed to BM-MDSC (untreated or infected, see above) supernatant either during the CD3/CD28 beads stimulation (24hrs, labelled ‘Exposed to supernatant during stimulation’) or after removal of the beads (72hrs, labelled ‘Exposed to supernatant after stimulation’). T cell proliferation was assessed through Cell Trace Violet dilution measurement via flow cytometry (A-C). Data show median values (n = 3 independent experiments, 2 to 5 replicate per experiment). Statistical significance: ANOVA, multiple comparison, Tukey’s test, ∗p < 0.05; ∗∗p < 0.01; ∗∗∗p < 0.001; ∗∗∗∗p < 0.0001. OT-1 CD8 T cell proliferation: MDSCs supernatants from mock-treated-(NT) or PAO1-infected cells were evaluated on the proliferation of OT-1 CD8 T cells stimulated with OVA peptide pulsed dendritic cells (D). Independently, T cells were also treated with supernatants of PAO1 (PAO1 WT only). Data show median values (n = 3 independent experiments, 1 to 2 replicate per experiment). Statistical significance: Kruskal-Wallis, Dunn’s multiple comparison, ∗p < 0.05; ∗∗p < 0.01; ∗∗∗p < 0.001; ∗∗∗∗p < 0.0001.

Even though we observed a reduced inhibitory effect when compared with the ‘full time’ exposition, there was inhibition of the proliferation in both conditions. This suggests that both compartments (synaptic and post-synaptic) may be targets of the inhibitory supernatants.

b) *Specific TCR stimulation*

In order to test more widely the inhibitory potential of infected MDSC supernatants, an antigen specific (OVA peptide) proliferation assay with isolated T cells from OT-1 mice was set up. Again, the supernatant from *P.a-*infected MDSC showed a strong inhibitory activity on CD3+CD8+ T cell proliferation (Fig. 7D).

### MDSC lung adoptive transfer rescues *P.a*-induced mice mortality

Since MDSC were shown above to have clear *in vitro* regulatory properties, we then tested their activity *in vivo*, in both prophylactic and therapeutic set-ups of *P.a* lung infection in mice.

In a preliminary experiment, we transferred MDSC through the oro-pharyngeal route and checked (with mismatched CD45.1 cells into CD45.2 C57/Bl6 receivers) that they did stay in the lung up to 24hrs (not shown).

With that information in hand, 2e6 ER-Hoxb8-MDSC (C57/Bl6 background) were then similarly transferred through the oro-pharyngeal route into male C57BL/6 mouse receivers either 4 hours before (in a prophylactic set-up (Fig. 8A, top)) or 16 hours after (in a therapeutic set-up (Fig 8A, bottom) lung PAO1 infection (1e8 CFU/mice, administered through the same route). Mice receiving MDSC had significantly lesser weight loss (Fig. 8B) and severity score (Fig. 8C). Mice survival was also very significantly improved in both MDSC receiver groups (H-4 or H+16), with 100% survival rate for the H+20 MDSC group.

**Figure 8.**
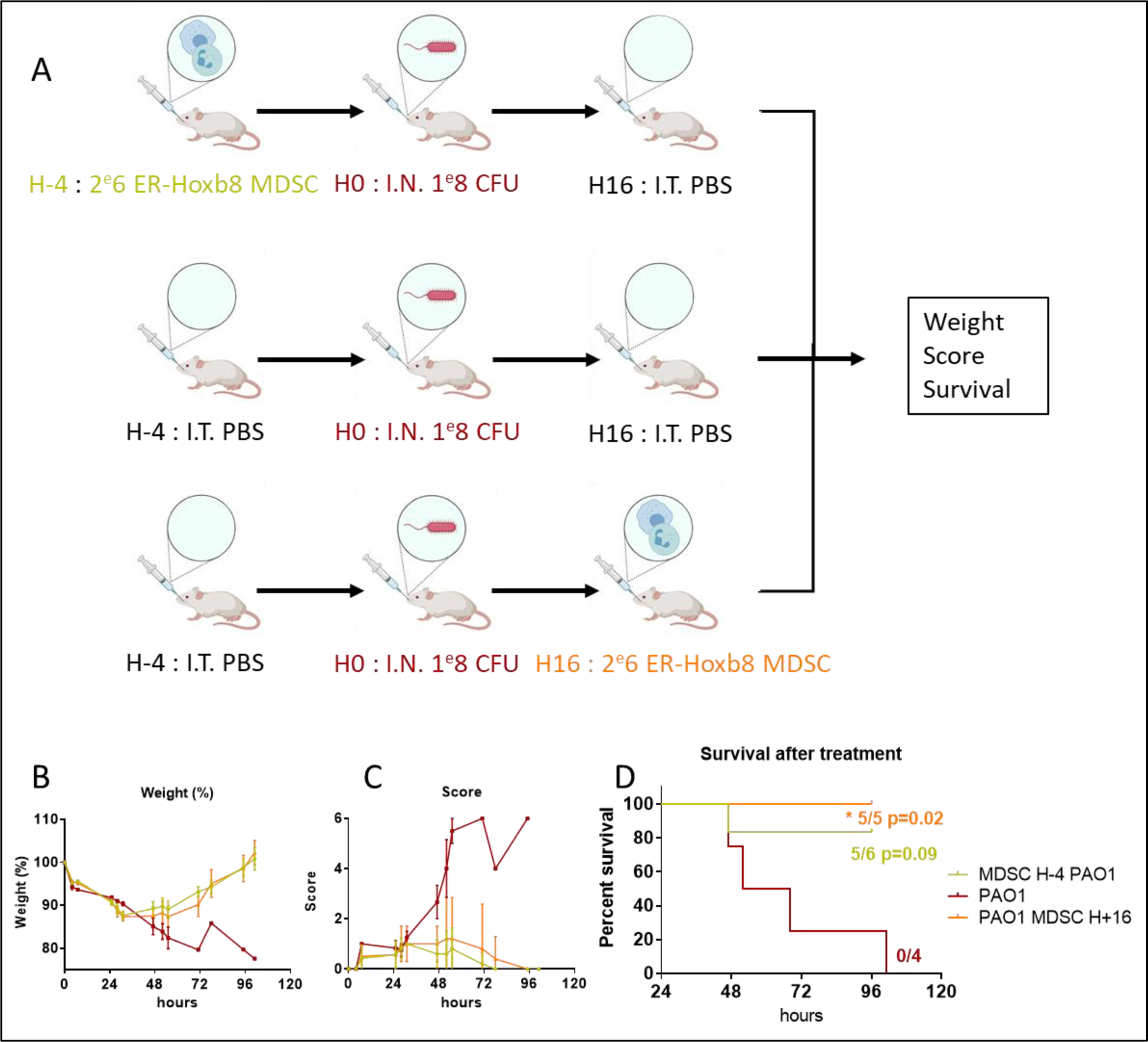
Lung adoptive transfer of ER-Hoxb8-MDSCs protects mice against lethal *Pseudomonas aeruginosa* (PAO1) infection. **(A).** 2e6 ER-Hoxb8-MDSCs were transferred into male C57BL/6 mouse receivers 4 hours before (A, top section, n=6) or 16 hours (A, bottom section, n=5) after PAO1 infection (1e8 CFU/mice). The control ‘PAO1 alone’ arm of the experiment (4hrs post PBS, 16hrs pre-PBS, n=4) is presented in A, middle section. Weight (B), score (C) and survival (D) were monitored. Survival was plotted with Kaplan-Meier curves, and statistical testing (against PAO1 alone) was performed with the log-rank (Mantel-Cox) test.

### PAO1 infected MDSC supernatant accelerate epithelial cells repair after scratch injury

Since 100% mice survival rate was obtained when MDSC were administered 20hrs after PAO1 infection, we hypothesized that MDSC may have a role in lung repair. This was further modeled *in vitro* using the murine CMT-2 alveolar cell line. After a mechanical scratch, we observed that healing in CMT-2 cells exposed to the supernatant from PAO1-infected MDSC was indeed accelerated compared to cells exposed to supernatant from mock-treated cells (Fig. 9).

**Figure 9.**
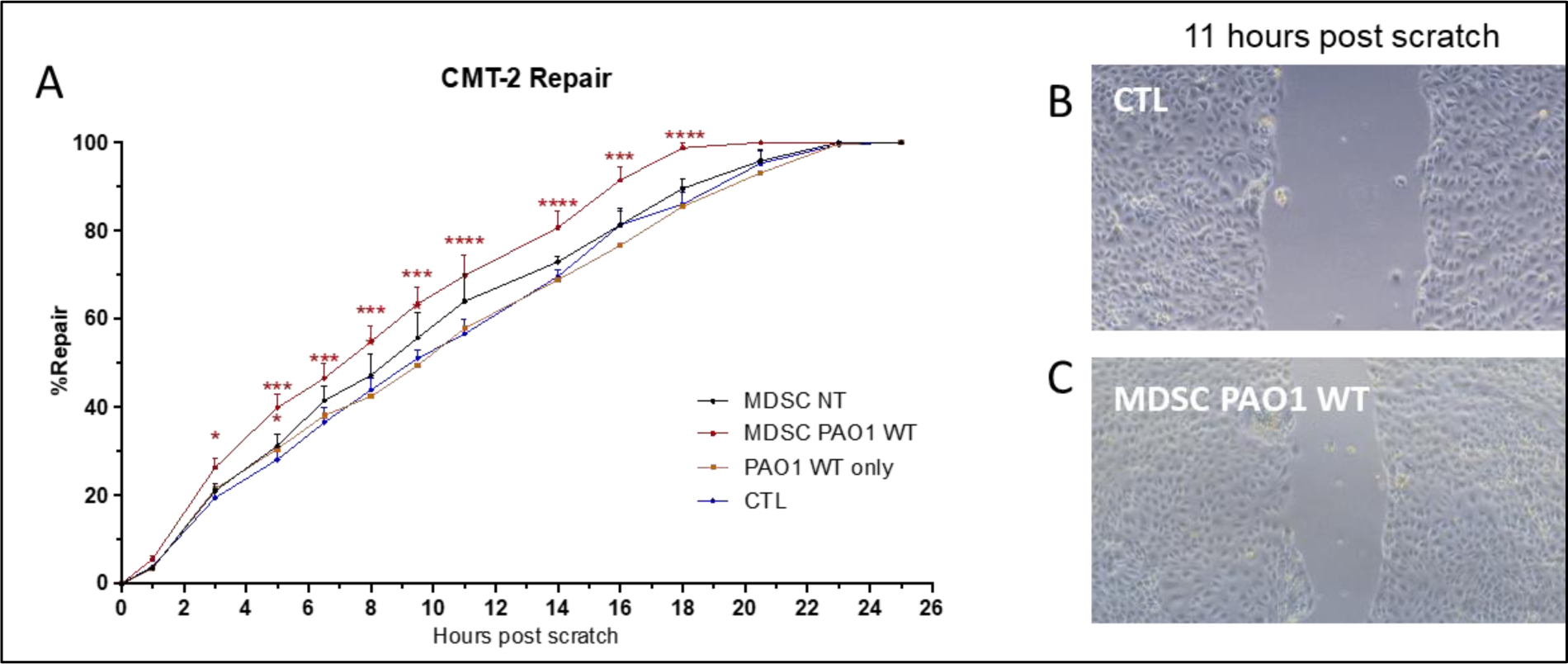
Supernatant of PAO1 infected MDSCs accelerate epithelial repair in vitro. In vitro differentiated BM-MDSCs were mock-(MDSC NT = non treated) or infected with PAO1 (MDSC PAO1 WT) for 6 hours at MOI 1. Supernatants were collected and filtered on 0.2µm filters. Repair inducing capacity of MDSC supernatants was evaluated in a CMT-2 alveolar cell line scratch assay. PAO1 cultures supernatants were also added independently (PAO1 WT only). CTL means CMT-2 cells exposed to unconditioned DMEM media. % of Repair (A) was measured by microscopy imaging (B-C) for 26 hours. Data show median ± SEM (n = 3 independent experiments). Statistical significance: Two Way ANOVA, multiple comparison ∗p < 0.05; ∗∗p < 0.01; ∗∗∗p < 0.001; ∗∗∗∗p < 0.0001.

## DISCUSSION

Previous studies have shown that anti-inflammatory/regulatory macrophages may be of interest in *P.a* infections for ameliorating lung function and overall survival^7–9^. Specifically, we showed that IL-6-genetically engineered macrophages provided a beneficial anti-inflammatory/reparative phenotype *in situ* in murine lungs, and protected mice against *P.a* infection^7^. Because IL-6 is an important factor for the differentiation of *bona fide* anti-inflammatory myeloid-derived suppressor cells (MDSC), we wished here to study their potential functions in this model. Indeed, while MDSC are ‘professional’ regulators well studied in cancer and autoimmune diseases, their function in infectious disease remains poorly understood^10^. For that purpose, we used here two types of murine MDSC, one derived from primary bone-marrow cells, as described by Lee at *al* in 2018^21^ and the other from the immortalized ER-Hoxb8 cell-line^22,27^.

The latter has been shown to be useful for DC, macrophage and neutrophil differentiation^22,28,29^ but until recently, no protocol had been reported to obtain MDSC from this cell line. Lately however, Allatar et al successfully obtained monocytic MDSC from these cells ^30^, using GM-CSF as a differentiating agent. Here, using an improved protocol (5-day exposition to both GM-CSF and IL-6, the same protocol used to obtain BM-MDSC), we obtained both monocytic and neutrophilic subsets of MDSC (Fig 1A-D), and showed that these cells could be expanded more efficiently than BM-MDSC (Fig 1E).

Regardless of their origin, we showed that when both BM-MDSC and ER-HoxB8 cells were infected with either the *P.a* laboratory strain PAO1, or *P.a* clinical strains, their supernatants strongly inhibited lymphocyte proliferation (Figs 2-3). This was at first puzzling since even though MDSC were known to secrete many effectors with immune suppressive capacity, a contact-dependent was thought crucial for MDSC function^31–33^. Interestingly, supernatants derived from infection of murine epithelial cells with PAO1, in the same conditions, showed no inhibitory activity (Fig 2 panels V-Y), demonstrating *P.a-* MDSC specificity. Mass spectrophotometric analysis showed, as expected, that the supernatants from the PAO1-MDSC incubation contained both bacterial- and host-derived proteins (not shown). Importantly, secretomes derived from PAO1 cultures *only* (at the same concentration) did not affect lymphocyte proliferation (e.g. Fig 2 panels S-U, Fig 3 all panels, Fig 4 panels J-K, etc..). Although it cannot be ruled out that *P.a* strains directly inhibits lymphocyte proliferation through the release of bacterial-derived mediators induced after the interaction with MDSC, a eukaryotic origin for this inhibitory factor seemed more likely. We showed, in set-ups involving a variety of bacterial mutants and experimental conditions that these mediators are derived from both M- and PMN-MDSC (Fig. 4), are heat- and proteinase K-resistant (i.e. likely not proteins, Fig. 6), and act downstream from the T cell synapse (Fig. 7).

We confirmed, by comparing ‘inhibitory supernatants’ (from PAO1 WT infections) with ‘non-inhibitory supernatants’ (from PAO1 d*fliC* infections) that typical inflammatory/anti-inflammatory cytokines levels were similar in both conditions (IL-1B, IL-6, IL-10, TGF-B ELISA, not shown.)

Given the described activity of lipid mediators (particularly PGE2) and extracellular vesicles as potential effectors of MDSC mediated suppression^34–36^, this was also tested here. However, in our set up, PGE2 concentration did not correlate with MDSC suppression, and inhibitors of COX-2 and cPLA2 did not affect the inhibitory activity of supernatants (not shown). Furthermore, the purified vesicular fraction of the supernatants did not induce T cell inhibition (not shown).

At the bacterial level, because flagellin has been shown to induce MDSC differentiation from PBMCs^19^, we set up to investigate whether flagellin or the flagella were also important in transmitting this inhibitory signal onto MDSC. Even though we showed that bacteria had to be alive (Fig 5A-C), motile (panels 5J-L), and bear a flagellum (panels 5D-F) to induce an inhibitory signal onto T cells via MDSC, this signal was shown to be independent of the TLR/Myd88 pathway (panels 5G-I), despite the description of the latter as being important for *P.a-*myeloid cells interaction^25,37–40^.

Indeed, it is more the mobility function of the flagella than the signaling property of its component flagellin which seemed important here. By allowing bacterial movement, the flagellum can promote contact between the cells and the pathogen and facilitate the contact with immune cells and the ensuing activation. This proposition is supported by previous work from Floyd et al, who showed that neutrophil extracellular trap release is activated by motile bacteria only, in a pathway independent of direct flagellin signaling^41^. A complementary possibility is that dFliC mutants may affect other virulence factors independently of motility. This hypothesis is sustained by a study from Suriyanarayanan et al who showed that phosphorylation mutants of the flagellin FliC protein affected the secretome of the type 2 secretion system (T2SS) of *P.a*^42^, a seemingly unconnected system.

Alternatively, other signaling pathways could be involved. For example, it is well known that upon phagocytosis, pathogen associated molecular patterns (PAMPs) can be recognized by cytosolic receptors such as NOD like receptors (NLRs). While we did not test their potential involvement here, they have been described by us and others to recognize *P.a.* related PAMPs^43^ and activate the inflammasomes, notably NLRC4^25,39,44,45^.

Regardless of the exact molecular mechanisms underlying the *ex-vivo* MDSC-derived inhibition of T cell proliferation, it remained to be tested whether this phenotype might be beneficial *in vivo.* Indeed, Rieber et al. showed that the MDSC proportion was increased in CF patients PBMCs infected with *P.a* and that this correlated with ameliorated lung function^19^. However, a strategy of MDSC adoptive transfer via the tail vein in mice previously showed no survival benefit^20^, but no information was given as to whether sufficient MDSC numbers reached the lung during infection to exert a biological effect. By contrast, we showed a beneficial effect of MDSC transferred through the oro-pharyngeal route, both in prophylactic and therapeutic set-ups (Fig 8). Although the mechanisms underlying this protective effect remain to be firmly established, it is possible that akin to what was modeled *in vitro*, *in vivo* transferred MDSC establish, in conjunction with *P.a,* an anti-inflammatory/reparative milieu in the lung propitious to accelerate survival. Although to our knowledge, this has not been previously specifically evaluated, this new concept is supported by our *in vitro* epithelium scratch assay whereby PAO1-infected MDSC supernatants accelerated wound closure.

In conclusion, we believe that the unique properties of MDSC make them attractive potential new therapeutic tools for patients with acute or chronic inflammatory diseases, where inflammation has to be kept in check. These cells could be added to the existing armamentarium for myeloid cell transplantation, currently including differentiated macrophages or stem cells^8,46–49^. Relatedly, recent innovative studies have indeed pointed towards the use of modified macrophages in alveolar proteinosis^50–52^, recently culminating in the first clinical trial using this methodology (ClinicalTrials.gov : NCT05761899)^53^.

## MATERIAL AND METHODS

### *In vivo* experiments

#### Animals

Seven- to ten-week-old male C57BL/6 WT and were purchased from Janvier Labs (Le Genest-Saint-Isle, France). Bone-marrows from TLR5 and Myd88 -/- were a gift from Dr. J.-C. Sirard (Institut Pasteur, Lille, France). Animals were kept in a specific pathogen-free facility under 12-h light/dark cycles, with free access to food and water. Procedures were approved by our local ethical committee and by the French Ministry of Education and Research (agreement number 28050).

#### MDSC adoptive transfer and PAO1 lung infection

Mice were either:

1. mock-instilled with PBS or instilled with ER-Hoxb8 MDSC (2x10^6 MDSC) and 4hrs later infected with 10^8 cfu of PAO1.
2. Infected with 10^8 cfu PAO1 and 16hrs later, re-instilled with either PBS or 2x10^6 ER-Hoxb8 MDSC.
3. Survival and weight were followed over time and behavior score (appearance and mobility observation combined with weight loss measurement) recorded.

MDSC instillations were performed through the oro-pharyngeal route after ketamine/xylazine anesthesia (100μL of ketamine 500 and xylazine 2% in 0.9% NaCl (10:10:80) intra peritoneal). Bacterial infections were done intranasally after isoflurane anesthesia.

### *Ex vivo* experiments

#### BM-MDSC differentiation

Bone marrow was extracted from mice femurs, and cells were washed with PBS and centrifuged at 1,400 rpm for 10 min (4°C). Pelleted cells were then re-suspended with ACK lysis buffer (Gibco) for 2 min at room temperature to lyse red blood cells. After another wash, cells were seeded in complete DMEM medium (10% FCS, 1,000 U/mL penicillin, 100 μg/mL streptomycin) containing 30 ng/mL of mouse granulocyte-macrophage colony-stimulating factor (GM-CSF) and Interleukin-6 (IL-6) (PeproTech). After 5 days, BM-MDSC were detached with Cell Dissociation Buffer (Gibco, 13151014), washed, and re-suspended in sterile PBS. For *in vitro* experiments, cells were seeded in 24-well culture plates (1500000 cells/well) in DMEM-Glutamax (10% FCS, 1,000 U/mL penicillin, 100 μg/mL streptomycin) for 12 h prior to stimulation or infection. For some experiments MDSC subset (PMN- and M-MDSC) were separated with the Miltenyi Biotech MDSC separation kit (Reference 130-094-538).

#### T cells proliferation assay

Spleens of WT C57BL/6 mice were harvested, processed, and homogenized into a single-cell suspension through mechanical disruption. Briefly, the spleens were grinded and filtered on a 40-μm sterile filter. Filters were washed with RPMI-Glutamax (10% FCS, 1,000 U/mL penicillin, 100 μg/mL streptomycin), and the recovered cells were centrifuged (2,000 rpm, 4°). Pelleted cells were re-suspended for 5 min at room temperature with 1 mL of ACK Lysis buffer to lyse red blood cells. Lysis reaction was stopped by adding 30 mL of sterile PBS 1×, and cells were centrifuged (2,000 rpm, 4°C). Pan T cells were then isolated with Miltenyi Biotech Pan T cell isolation kit II (Reference 130-095-130) according to manufacturer’s protocol.

Isolated T cells were re-suspended at 1 × 10^6/mL in 37°C pre-warmed PBS and incubated for 20 min at 37°C with 2μM Cell Trace Violet (CTV) (Invitrogen). Cells were washed in RPMI-Glutamax (10% FCS, 1,000 U/mL penicillin, 100 μg/mL streptomycin). The CTV-labeled cells were seeded in a 96-well culture plate (1×10^5 cells/well) in R20 medium (20% FCS, 1,000 U/mL penicillin, 100 μg/mL streptomycin, 50µM β-mercaptoethanol, 1X Non-Essential Amino-Acids, 1X Sodium pyruvate) supplemented with 20ng/mL of mouse rIL-2. 2.5 μL of Dynabeads Mouse T-Activator CD3/CD28 (Gibco Life Technologies) were added in each well except for a control well (negative control). 100 μL of PBS (control), DMEM (control), MDSC supernatant was then added into the wells, and cells were left in culture for 72 h. Cells were then harvested, stained for CD3, CD4, CD8 and washed with FACS buffer (PBS 1% BSA, 0.5mM EDTA), and CTV intensity was analyzed by flow cytometry. In a subset experiment, T cells were exposed to Dynabeads for 24 hours. After stimulation, well contents were harvested, beads were removed thanks to a magnetic bench, cells were washed and put in culture in R20 medium, with or without MDSC supernatant, for 3 days.

In independent experiments, supernatants from mock- or PAO1-infected MDSC were incubated with T cells from OT-1 mice spleens (a kind gift from Dr. L. Saveanu) previously co-cultured with dendritic cells (MutuDC cell line) pulsed with 1nM OVA during 2 hrs.

### *In vitro* Experiments

#### Cell lines

##### ER-Hoxb8 culture and ER-Hoxb8-MDSC Differentiation

The ER-HoxB8 murine cell line^22^ (C57/Bl6 background) was a kind gift from Dr D. B. Sykes (Center for Regenerative Medicine, Massachusetts General Hospital, Boston, MA, USA).

ER-HoxB8 cells were cultured in RPMI-Glutamax (10% FCS, 1,000 U/mL penicillin, 100 μg/mL streptomycin) supplemented with 30 μM β-mercaptoethanol, 1% CHO-SCF conditioned media and 1µM β-estradiol in 6 well plates at 500.000 to 3.000.000 cells/well. ER-HoxB8 cells were split 1:5 every 48-72 h with the addition of a fresh culture medium.

For differentiation in MDSC, β-estradiol was removed by washing cells 3 times in PBS. 500.000 cells were seeded in 10cm non adherent Petri dishes with complete DMEM medium (10% FCS, 1,000 U/mL penicillin, 100 μg/mL streptomycin) containing 30 ng/mL of mouse granulocyte-macrophage colony-stimulating factor (GM-CSF) and Interleukin-6 (IL-6) (PeproTech) and 4% CHO-SCF conditioned media. After 5 days, ER-HoxB8 MDSC were detached with Versene 1X (Gibco), washed, and re-suspended in sterile PBS. For *in vitro* experiments, cells were seeded in 24-well culture plates (1500,000 cells/well) in DMEM-Glutamax (10% FCS, 1,000 U/mL penicillin, 100 μg/mL streptomycin) for 12 h prior to stimulation or infection. For *in vivo* adoptive transfer experiments, MDSC density was adjusted to 2x10^6 cells/50 μL in sterile PBS before oro-pharyngeal instillation.

##### CMT-2 and DJS-2

Mouse alveolar epithelial cell line CMT-2 (CMT64/61, European Collection of Authenticated Cell Cultures, ECACC) and mouse bronchial epithelial cell line (Club Cells) a kind gift from Dr. DeMayo (Baylor College of Medicine, Houston, TX, USA), were cultured in DMEM-Glutamax (10% FCS, 1,000 U/mL penicillin, 100 μg/mL streptomycin). Cells were placed for 12 h (37°C, 5% CO2) in 24-well Corning Costar culture plates (750,000 cells/well) prior to scratch assays.

#### Scratch assay

Cell monolayers were injured mechanically with BioTek Auto Scratch Wound Making Tool (https://www.agilent.com/en/product/cell-analysis/cell-imaging-microscopy/imager-multimode-reader-peripherals/biotek-autoscratch-accessory-supplies-1623134). Afterwards, cells were washed with their culture medium to remove detached cells and were treated with DMEM (control) or MDSC supernatants. Photographs of the wounds were taken using an inverted microscope with an X10 objective at different time points after scratch. Images were analyzed using ImageJ software (https://imagej.nih.gov/ij/index.html) and Wound Healing Tool updated plug-in (https://github.com/AlejandraArnedo/Wound-healing-size-tool/wiki) to measure areas of the wounds and mean wound closure (% of the area at t = 0 h) was calculated.

### Bacteria

#### PAO1

PAO1 strain (ATCC 15692) was kept in freezing medium (50% Luria broth [LB], 50% glycerol) and stored at −80°C until use. Before infection experiments, PAO1 strain was grown overnight in LB in a rotating incubator (200 rpm, 37°C). The bacterial suspension was then diluted in serum- and antibiotic-free RPMI medium, and the optical density (OD) was measured at 600 nm every 1 h until logarithmic growth phase was reached (0.1 < OD < 0.3; an OD of 0.1 is equivalent to a bacterial concentration of 7.73 × 10^7 CFU/mL). PAO1 WT, d*fliC* and d*pscN* were kind gifts from Dr. Romé Voulhoux (LCB-UMR7283, CNRS, Aix Marseille Université) and was used under the same protocol as PAO1.

#### Clinical strains

Clinical isolates were kind gifts from Dr. C. Llanes and P. Plésiat (Laboratoire de Bactériologie UMR CNRS 6249 ; UFR SMP, Besançon, France) and used as previously described^54^.

##### Streptococcus Pneumoniae

*S. pneumoniae* (clinical isolate E1586, gift from Dr. J-C. Sirard) was stored in −80°C in freezing medium (50% Todd Hewitt Yeast THYB/50% Glycerol) and used as PAO1 in infection experiments.

#### Cell infection assays

MDSC (or their fractions) were either mock-treated or infected with PAO1WT and mutants for 6 h at MOI=1.

Supernatants were collected and filtered on 0.2µm filters before functional assays (T cell proliferation, scratch assays) or ELISA analysis were performed. In subset experiments, PAO1 supernatants alone were produced by adding a quantity of PAO1 equal to infection inoculum (MOI = 1, i.e. 1,5e6 CFU) in a separated well and let 6 hours in culture (steady plate, 37°C, 5% CO_2_). After 6 hours, media was harvested, filtered and added to T cells proliferation assays.

Cells were lysed with RNA lysis buffer +50uM β-Mercaptoéthanol or collected for flow cytometry analysis.

Epithelial cell lines infection experiments were performed under the same protocol.

### RNA Extraction, qPCR, ELISAs

#### RNA extraction, reverse transcription, and qPCR

Cellular RNA isolation was performed with the PureLink RNA Mini Kit (12183018A, Ambion, Life Technologies), following the manufacturer’s instructions. 500ng of total RNA was treated with DNAse (Roche), and retrotranscribed to cDNA using random hexamers (Roche) and M-MLV reverse transcriptase (Promega). Real-time PCR was performed in a QuantStudio 6 Flex PCR System (Applied Biosystems) using the Fast SYBR Green Master Mix (Applied Biosystems). Results are expressed as dCT = CT gene of interest-CT housekeeping gene (18S). qPCR primers are reported in Table 1.

#### ELISA

All ELISA kits were purchased to R&D Biosystems (DuoSet ELISA) and performed according to manufacturer’s protocol.

### Flow cytometry

#### Classical analysis

MDSC were detached from culture plates with Cell Dissociation Buffer (Gibco, 13151014), followed by a PBS wash, for completion of detachment. Cells were then first incubated (10 min, 4°C) with a cocktail of a viability dye and Fc Block antibody, washed (2,000 rpm, 5 min) with FACS buffer (PBS-2% FCS) and then incubated (30 min, 4°C) with a cocktail of cell surface conjugated antibodies. Finally, cells were further washed, and the pellets were re-suspended in FACS buffer for data acquisition. Data were acquired the same day with a LSRFortessa cytometer (BD Biosciences) with BD FACSDiva software and analyzed with FlowJo (Tree Star, Ashland, OR, USA). The antibodies used for FACS analysis are listed in Table 2.

### Statistical analysis

Data were analyzed with GraphPad Prism Software 6.05. Statistical analysis was performed with either a non-parametric test (Kruskal-Wallis and Wilcoxon) or one-way or two-way ANOVA followed by the appropriate multi-comparison post hoc Tukey’s test.

Survival curves in murine model experiments were plotted with Kaplan-Meier curves, and statistical testing was performed with the log-rank (Mantel-Cox) test.

Differences were considered statistically significant when p was <0.05 and are labeled as follows: ∗p < 0.05; ∗∗p < 0.01; ∗∗∗p < 0.001; ∗∗∗∗p < 0.0001.

## Acknowledgments

We wish to thank “Vaincre la Mucoviscidose” for recurrent support, and for the award of a PhD grant to MBB (RF20200502676 and RF20230503234). We are also thankful to Drs. J-C Sirard and L. Van Maele (Institut Pasteur, Lille, France), Dr. L. Saveanu (Centre de Recherche sur l’Inflammation, Paris, France), Dr. R. Voulhoux (LCB-UMR7283, CNRS, Aix Marseille Université), Dr. DeMayo (Baylor College of Medicine, Houston, TX, USA), Dr. D. B. Sykes (Center for Regenerative Medicine, Massachusetts General Hospital, Boston, MA, USA), Dr. C. Llanes and Dr. P. Plésiat (Laboratoire de Bactériologie UMR CNRS 6249; UFR SMP, Besançon, France) for all the kind gifts described above and Mrs G. Ball (LCB-UMR7283, CNRS, Aix Marseille Université) for generating *Pseudomonas* mutants.

## MATERIAL AND METHODS TABLES

**Tableau 1.**
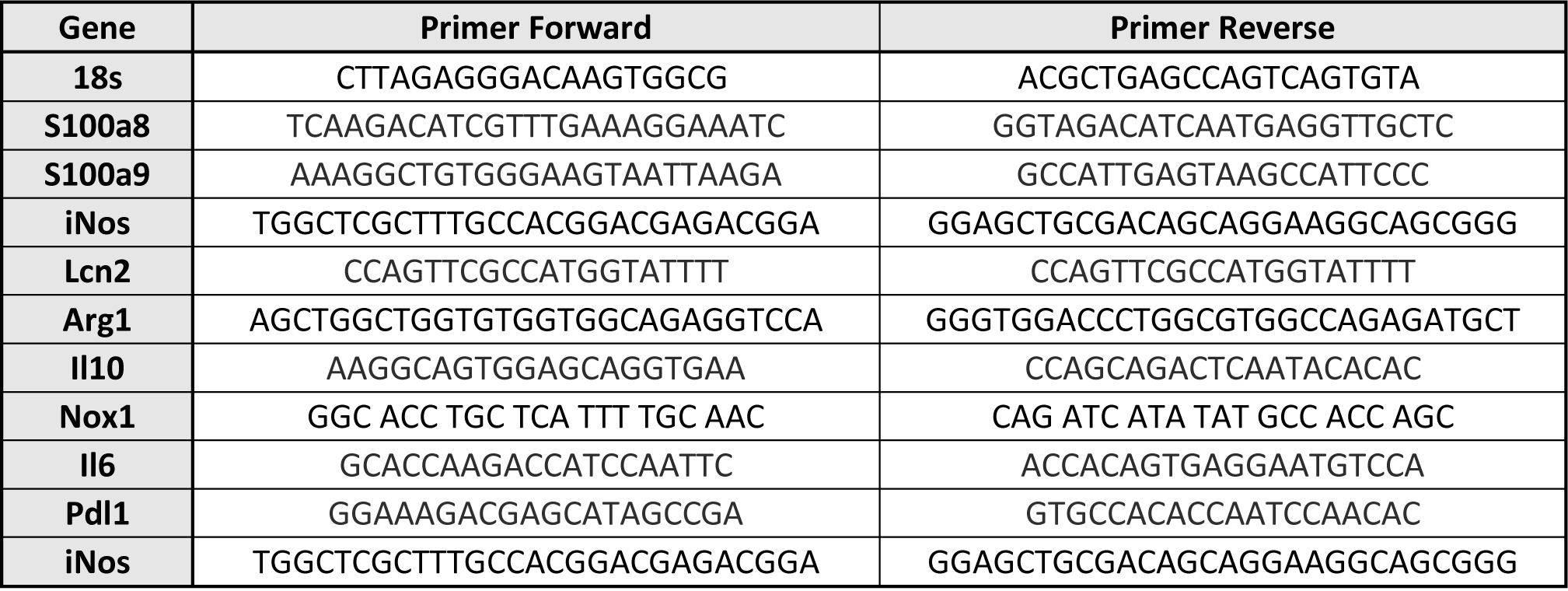
qPCR Primers.

**Tableau 2.**
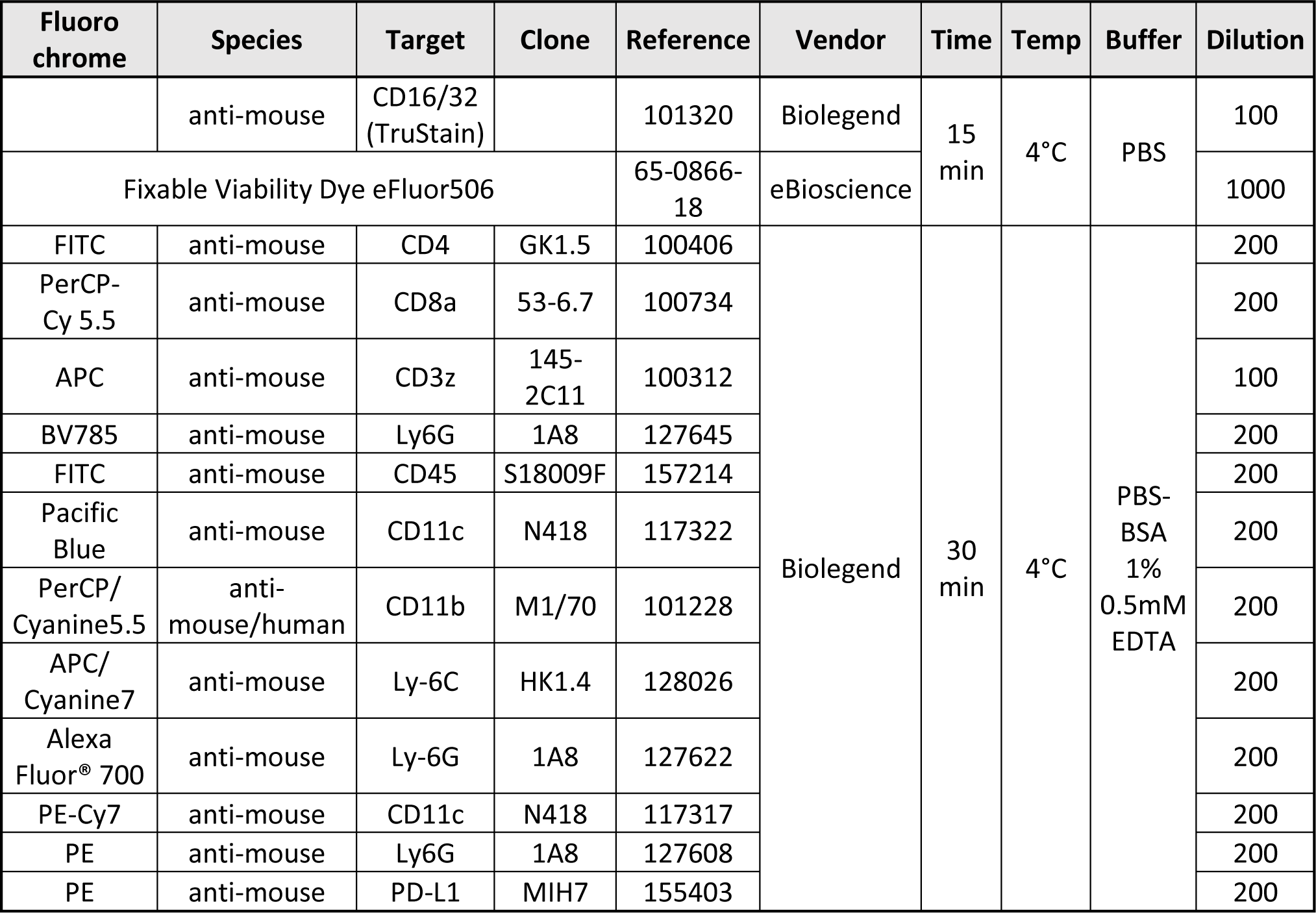
Flow cytometry antibodies.

**Tableau 3.** *In vivo* scoring.

Tableau 2 - Flow cytometry antibodies
| Appearance |  |
| --- | --- |
| Normal | 0 |
| Poorly groomed appearance | 1 |
| Dull coat, slight nasal/ocular discharge | 2 |
| Ruffled coat/hunched abdomen or protruding abdomen/hunched back | 4 |
| Behavior |  |
| Normal/exploratory | 0 |
| Labored breathing/hunched posture | 1 |
| Apathy/total indifference to the environment/inability to move | 4 |
| Weight |  |
| Weight loss between 10-15% | 1 |
| Weight loss between 15-20% | 2 |
| Weight loss between 20-25% within 24 hours | 4 |
| Weight loss between 20-25% over 24 hours | 5 |

## Notes

### Competing Interest Statement

The authors have declared no competing interest.

